# Opioid mortality following implementation of medical marijuana programs (1999-2017) in the United States

**DOI:** 10.1101/670059

**Authors:** Daniel E. Kaufman, Asawer M. Nihal, Janan D. Leppo, Kelly M. Staples, Kenneth L. McCall, Brian J. Piper

**Affiliations:** Geisinger Commonwealth School of Medicine, Scranton, PA, USA; University of New England, Portland, ME USA; Center for Pharmacy Innovation and Outcomes, Forty Fort, PA, USA

**Author notes:** Brian J. Piper, PhD MS, Department of Medical Education, Geisinger Commonwealth School of Medicine, Scranton, PA 18411, or.

**Keywords:** Medical marijuana, opioids, overdose, mortality, cannabis

## Abstract

The United States is in the midst of an opioid overdose epidemic. A prior report using the Center for Disease Control’s Wide-ranging Online Data for Epidemiologic Research (WONDER) database discovered that opioid overdoses decreased by 24.8% from 1999 to 2010 in states with medical cannabis (MC+) relative to those without (MC−). The present study evaluated any differences following MC legislation on WONDER reported opioid overdoses, corrected for population, from 1999 to 2017 using an interrupted time series. Overdoses were significantly higher in MC+ states from 2012-2017. The slope of opioid overdose deaths over time increased significantly post-implementation in states without MC (3-years Pre = 0.1 ± 0.1, 3-years Post = 0.7 ± 0.2, *t*(16) = 2.88, *p* ≤ .011). Overdose deaths showed a non-significant elevation in states with MC (Pre = 1.3 ± 0.3, Post = 2.8 ± 0.8, *t*(11) = 2.01, *p* = .069). Post-legalization slopes were significantly higher in MC+ than MC− (*t*(11.95) = 2.70, *p* < .05). Overall, any impact of medical cannabis laws on opioid overdoses appears modest. There are other confounds (e.g. death determination reporting quality) which differ non-randomly among states and are non-trivial to account for in ecological investigations of cannabis policy. Alternatively, the potency of fentanyl analogues may obscure any protective effects of MC against illicit opioid harms.

## Introduction

The US continues to struggle to reverse an opioid overdose crisis. Overdoses involving any opioid increased from eight-thousand in 1999 to approximately nineteen-thousand in 2007 and to almost forty-eight-thousand in 2017 (NIDA, 2019). Overdose data should be interpreted cautiously however as states differ widely in use of medical, or even non-medical, personnel or analytical chemistry procedures to complete death determinations. The number of overdoses where the substance involved was unspecified on the death certificate ranged from 0% in Washington DC to 51% in Pennsylvania (Buchanich et al. 2018). The Centers for Disease Control and Prevention (CDC) used a three-tier system (excellent, good, or other/less than good) to classify overdose death determinations with twenty-two states falling into the latter designation (Rudd et al. 2016). Interestingly, prescription opioid use, a measure with much more homogenous data collection, peaked in 2011 (Piper et al. 2018) and has undergone pronounced reductions for most agents, with the exception of buprenorphine (Collins et al. 2019). Buprenorphine availability is associated with decreased opioid overdoses (Schwartz et al. 2013). States that expanded Medicaid saw a seventy percent increase in buprenorphine prescriptions (Wen et al. 2017).

Several lines of evidence, albeit mostly from non-randomized and non-blind research designs, are suggestive of the potential for medical cannabis (MC) to attenuate opioid use or misuse. The potency of morphine on the rodent tail flick response to an aversive thermal stimuli was greatly enhanced by tetrahydrocannabinol (THC, Smith et al. 1998). Human trials have supported the ability of THC to augment the pain reducing effects of morphine and oxycodone (Abrams et al. 2011; Cooper et al. 2018). Three-quarters of dispensary members reported a reduction in their use of opioids after starting MC (Piper et al. 2017). The 2017 National Academy of Sciences report stated that evidence was conclusive that cannabis reduces chronic pain in adults, but the magnitude of effect was modest (National Academy of Sciences, 2017). Similarly, examination of a prescription drug monitoring program records revealed that patients were seventeen-fold more likely to stop use of all controlled substances after starting MC (Stith et al. 2018). States that legalized MC had lower expenditures for prescription medications in Medicare (Bradford & Bradford, 2016) and Medicaid (Bradford & Bradford, 2017, although see Ozluk, 2017). One of the most impactful studies was on opioid analgesic and heroin overdoses from 1999 – 2010 which made two discoveries. First, states that legalized MC had more opioid overdoses relative to those that did not. Second, opioid overdoses declined following MC implementation relative to those states without MC (Bachhuber et al. 2014). Further examination identified pre-existing state differences in opioid dependence hospitalizations and reductions associated with MC implementation (Shi, 2017). Findings like these (Lucas, 2017) may have been influential in for eight states (Colorado, Illinois, Missouri, Nevada, New Jersey, New Mexico, New York, and Pennsylvania) including opioid misuse as a qualifying condition for medical cannabis (NORML, 2019).

The objective of this report was to provide an update to Bachhuber et al. 2014 and examine opioid overdose mortality with the inclusion of additional states that have approved MC.

## Methods

### Procedures

The opioid overdose mortality rate from 1999-2017 in each state was extracted from the Centers for Disease Control and Prevention’s Wide-ranging Online Data for Epidemiologic Research (WONDER) database (https://wonder.cdc.gov/). The data was based on death certificates for US residents. Opioid overdose deaths were defined using the *International Statistical Classification of Diseases, 10^th^ revision* codes: X40-44, X60-64, X85, Y10-Y14 (Bachhuber et al. 2014). Only data that was coded for opioids (T40.0-T40.4) was used (Powell & Pacula, 2018). This included opium, heroin, other opioids, methadone, and other synthetic opioids.

An interrupted time series examined any trends around the time of states’ medical marijuana program implementation. Opioid deaths per 100,000 population were obtained. We defined the start dates of medical marijuana programs as the year the state began medical marijuana sales. This data was found on either state’s medical marijuana program website, or from local newspaper articles detailing the start of medical marijuana sales. Only Arizona (implemented in 2012), Connecticut (2014), Delaware (2015), District of Columbia (2013), Illinois (2015), Maine (2011), Massachusetts (2013), Minnesota (2015), New Jersey (2012), New Mexico (2007), Rhode Island (2013), and Vermont (2004) had start dates within the year range of 1999-2017, with three years of data available both pre- and post-medical marijuana program implementation. The states without medical cannabis laws (Alabama, Georgia, Idaho, Indiana, Iowa, Kansas, Kentucky, Louisiana, Mississippi, Nebraska, North Carolina, South Carolina, South Dakota, Tennessee, Texas, Virginia, Wisconsin, and Wyoming), irrespective of cannabidiol only, served as a comparison group. Procedures were deemed exempt by the IRB of the University of New England.

### Data analysis

Statistical analysis was conducted using Systat software, v13.1. Opioid overdoses were compared for states with versus without MC (+ versus −). If the assumption of homogeneity of variance was not met (*p* < .10), a separate variance t-test was completed. Slopes for opioid overdoses from three years before the implementation of medical cannabis (pre), and three years after (post, the year of implementation was excluded as a transitional year) were calculated with GraphPad Prism 8. The mean medical marijuana program implementation dates (2012) was used as the interruption point for the comparison group.

Two potential confounds were also examined. A two (MC− vs MC+) by two (Medicaid expansion by 2014 or 12/2017) chi-square and a t-test on the CDC’s three-tiered classification system (Rudd et al. 2016) for the quality of death certificate reporting (2 = very good/excellent, 1 = good, 0 = less than good based on completeness and consistency over time, described further in the Supplemental Figure 1 caption) were completed. Variability was expressed as the SEM. A *p* value less than 0.05 was considered statistically significant. Heat maps were created using Excel.

**Figure 1.**
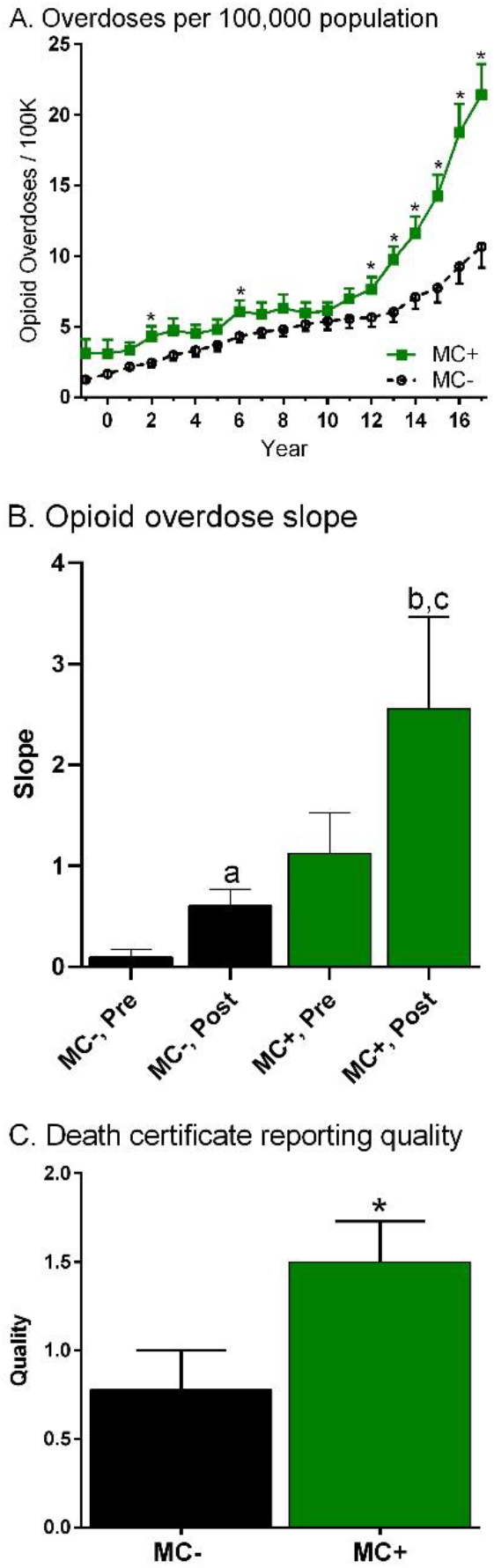
Opioid overdoses in the US as reported by the CDC’s Wide-ranging Online Data for Epidemiologic Research from 1999 to 2017 as a function of presence (+) or absence (−) of state medical cannabis (MC) law (A, * *p* < .05 versus MC−). Opioid overdose slopes three-years before (Pre) and three-years after (Post) state MC implementation (B, ^a^*p* < .05 versus MC−; ^b^*p* = .07 versus MC+ Pre, ^c^*p* < .05 versus MC− Post). Death certification reporting quality (C, Excellent: 2, Good: 1, Less than good: 0; \**p* < .05).

## Results

Figure 1A shows that overdoses had pronounced elevations over time and were generally higher among MC+ states. Significant differences in opioid overdoses per 100,000 population were identified in ’02, ’06, and ’13 to ’17 between MC+ and MC− state.

Figure 1B illustrates that the linear-regression slope of opioid overdose over time increased significantly in states without medical marijuana (Pre = 0.10 ± 0.08, Post = 0.61 ± 0.16, *t*(16) = 2.88, *p* < .05). The change in states with medical marijuana did not fulfill conventional statistical thresholds (Pre = 1.26 ± 0.40, Post = 2.77 ± 0.91, *t*(11) = 2.01, *p* = .069). The post-legalization slope was significantly higher in MC+ than MC− (*t*(11.95) = 2.70, *p* < .05) states.

Two MC− (11.1%) as compared to eleven MC+ (91.7%) states expanded Medicaid by 2014 (χ^2^(1) = 19.03, *p* < .0005). The same general pattern was noted when the expansion date was set at 2017 (MC− = 22.2%, MC+ = 91.7%, χ^2^(1) = 13.89, *p* < .0005). The quality of overdose determinations was twice as large in MC+ states (*t*(28) = 2.18, *p* < .05, Figure 1C).

## Discussion

This report re-examined and extended upon the seminal report of Bachhuber et al. 2014. The findings may be considered a partial replication. Their result that states with MC from 1999-2010 had elevated opioid mortality, relative to those without MC, was confirmed with the inclusion of seven additional years (Figure 1A). Their finding of opioid overdoses decreasing in the years subsequent to adopting MC as compared to states that did not, was not replicated. In fact, states that previously adopted MC had a significantly greater overdose slopes than those that did not (Figure 1B). The preclinical (Smith et al. 1998) and epidemiological evidence base (Bradford & Bradford, 2017, 2018; National Academy of Sciences, 2017; Stith et al. 2018) indicates that it is plausible that MC could decrease opioid use with its synergistic anti-nociceptive properties (Abrams et al. 2011; Cooper et al. 2018) and therefore limit misuse and overdose potential. However, investigations like Bachhuber et al. 2014 assume that overdose determinations are consistently made by medical examiners and coroners across the US (Supplemental Figure 1). At the very least, any differences should be random and not systematically different based on MC laws. An additional key finding was that this assumption was not warranted (Figure 1C) in that states without MC had lower overdose death determination reporting quality using the CDC criteria (Rudd et al. 2016).

The continued pronounced elevations in opioid overdoses, typically involving poly-substances (Simpson et al. 2019) is concerning. These findings, although largely contradictory to Bachhuber et al. 2014), should not be used to discount that the risk to benefit ratio may still favor MC as another tool that may be endorsed by some chronic pain patients, especially relative to the overdose potential of prescription opioids. As others have recently noted (Shover et al. 2019), the ecological research design where the unit of analysis is the state might not be best-suited to address this question. State-level MC systems, either established through voter initiative or through their state legislation, may differ on a wide-variety of socio-political characteristics. For example, Medicaid expansion was much more common among MC+ states. The additional resources provided by Medicaid could have provided for additional buprenorphine (Wen et al. 2017) which initially decreased opioid overdoses (Schwartz et al. 2013). Later, the potency of fentanyl and many fentanyl analogues (Simpson et al. 2019) could overcome any cannabis substitution effect (Piper et al. 2017). The increase in China white variety of heroin which is more common in the eastern US (Mars et al. 2016) and is more susceptible to contamination with white fentanyl than black tar heroin likely also contributes to regional differences in the recent overdose surge. This may be crucial because the majority (83.3%) of the MC+ states were east of the Mississippi.

Some caveats and future areas of study should be noted. This report was based on US overdoses as reported by the CDC. Five states had more than one-third of their death certificates where the substance involved was unspecified (Buchanich et al. 2018). The non-uniformity of autopsy procedures (e.g. When did each state, county, or city start testing for fentanyl? Were screens or the more expensive confirmatory testing employed? How many of the two-hundred fentanyl analogues were tested in each municipality and when?) is a substantial challenge to research, and barrier for empirically informed public policy which utilizes this information. Although this would limit the power to detect differences, future research might consider focusing only on the subset of areas where the death determination procedures have been consistently high (e.g. Simpson et al. 2019). This study investigated MC and may not generalize to recreational marijuana laws.

## Conclusion

New empirically grounded solutions to reverse the continued escalation in opioid overdoses are urgently needed. This study tested whether the protective effects identified by Bachhuber et al. 2014 of MC against opioid overdoses could be repeated with the addition of more data. States with MC had increased, not decreased as would be predicted based on Bachhuber et al. 2014, overdose slopes. An impediment to opioid overdose research is that overdose determinations procedures are not regionally or temporally uniform. States with MC had significantly higher quality of death certificate reporting. Additional investigations using other research designs and dependent measures are ongoing to further our understanding of the population health benefits, and risks, of MC.

## Supporting information

Overdoses by state

## Acknowledgements

The authors would like the thank Elizabeth Kuchinski, MPH for help and guidance throughout the research process. Amalie Kropp Lopez, MBS provided feedback on an earlier version of this manuscript.

---

**Supplemental Figure 1.**
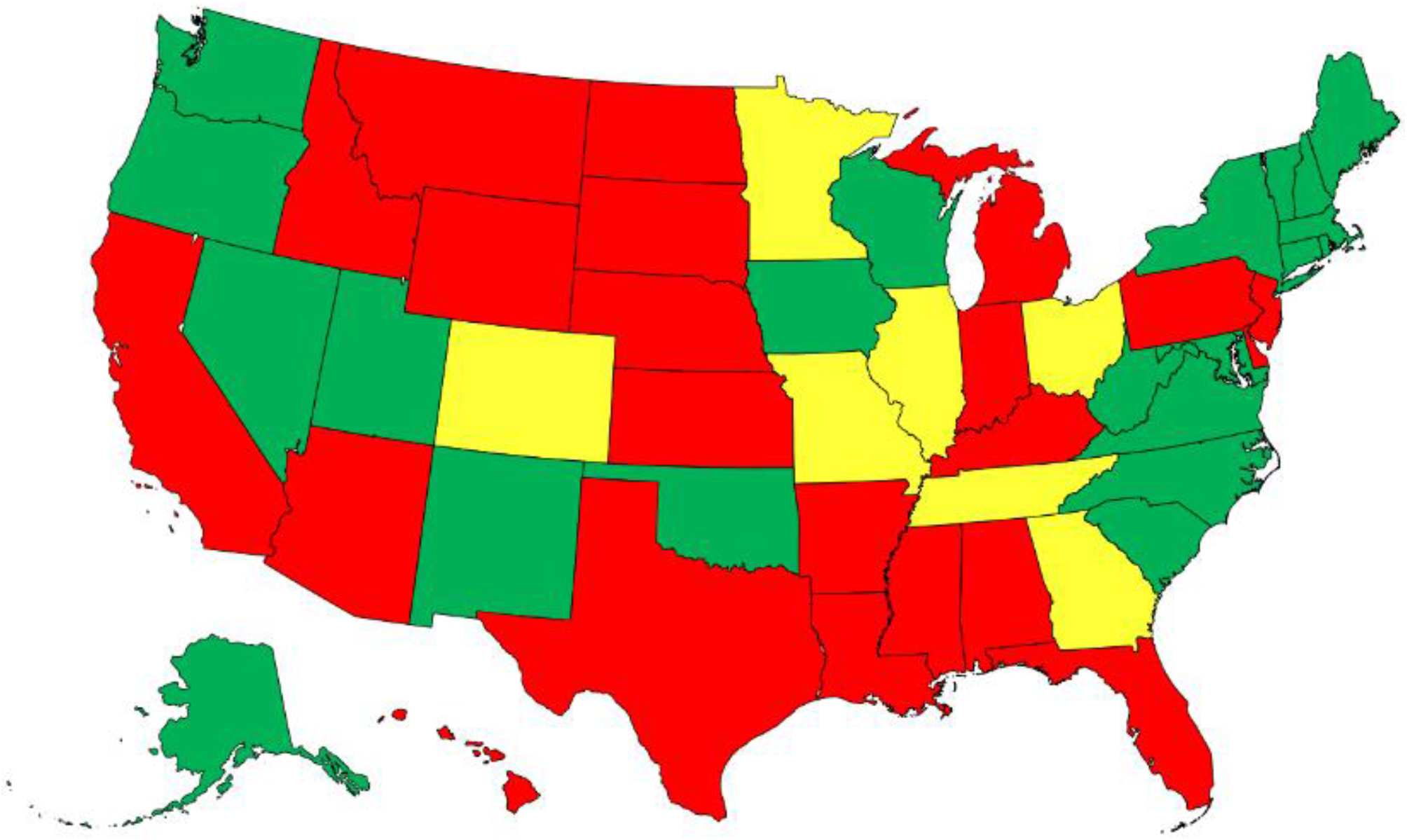
Heat map of the US by overdose death certificate reporting quality. Excellent was defined as ≥ 90% of drug overdose death certificates mention at least one specific drug in 2014, with the change in percentage of drug overdose deaths mentioning at least one specific drug differing by <10 percentage points from 2014 to 2015. Good was defined as 80% to <90% of drug overdose death certificates mentioning ≥ one specific drug in 2014, with the change in the percentage of drug overdose deaths mentioning ≥ one specific drug differing by <10 percentage points from 2014 to 2015. The remaining states were classified as less than good. Additional information may be found in Rudd et al. *Morbidity & Mortality Weekly Report* 2016; 65:1445-52.

